# Human lung γδ T cells maintain functionality during inflammatory lung disease

**DOI:** 10.64898/2026.04.23.720435

**Authors:** Alexis Taber, Marie Frutoso, Nicole Potchen, Amanda L. Koehne, Chelsea Schmitz, Eric D. Morrell, Martin Prlic, Shelton W. Wright

**Author notes:** address correspondence to: Shelton Wright, MD MS, Martin Prlic, PhD.

## Abstract

γδ T cells provide mucosal defense against infection while also contributing to tissue repair. However, data regarding the effect of the human lung environment on γδ T cell functionality remains limited. To address whether lung inflammation impacts γδ T cell functionality, we analyzed lung and matched hilar lymph node (LN) tissue from deceased donors and patients with interstitial lung disease (ILD). We performed high-parameter spectral flow cytometry to examine the expression pattern of phenotypic biomarkers and assess ex vivo function. We identified lung-specific enrichment of γδ T cells with an effector memory phenotype relative to matched regional LN. We then used an ex vivo stimulation approach to interrogate the capacity to protect against infection (granzyme B [GzmB], interferon-γ [IFNγ] and tumor necrosis factor [TNFα]) and promote epithelial cell proliferation (amphiregulin [AREG]). We found that γδ T cells in lung and LN from deceased donors had similar functional properties. While γδ T cell populations from ILD lungs largely maintained cytokine production capacity, expression was diminished relative to LN counterparts. Importantly, lung γδ T cells maintained polyfunctional GzmB, IFNγ and TNFα expression across cohorts. Overall, we report human lung γδ T cells are regionally distinct with conserved functionality in a fibrotic environment.

## Introduction

The lung, with millions of alveoli and an extensive airway network, represents a large and unique epithelial environment. This complex organ is supported, in part, by a robust lymphatic system, including hilar lymph nodes [1]. γδ T cells are an innate-like T cell subset capable of recognizing microbially-derived products, including phosphoantigens [2]. In mice, γδ T cells have been implicated as key immunologic players during lung inflammation, including regulating initial immune responses as well as initiating tissue repair [3]. Early in human development, this lymphocyte subset expresses a γδ T cell receptor (TCR) and can proliferate to various tissues, particularly in the mucosa, as well as their associated lymph nodes [4]. While they represent a relatively small lymphocyte subset, γδ T cells can recognize and respond to a variety of stimuli, including those beyond traditional MHC-peptides [5]. After activation, γδ T cells can express a variety of effector molecules, including IFNγ, TNFα, and AREG, which affect host immune responses as well as tissue repair [6–8]. Although many γδ T cell functions are conserved across species, substantial differences exist between γδ T cells derived from humans and mice, where they are most frequently studied. In humans, γδ T cell subsets are often defined by their TCRδ chain, resulting in two primary subsets: Vδ1 and Vδ2 [9,10]. The Vδ2 γδ T cell subset is most commonly studied in both disease and health, likely in part due to its enrichment in the peripheral blood. By contrast, murine γδ T cells are characterized by a multitude of relatively tissue-specific subsets, defined typically by their TCRγ chain [10,11]. Murine γδ T cells are also frequently categorized by their ability to express either IL-17 or IFNγ, eliciting specific effector functions, though how these functional phenotypes extend to human tissues remains any area of investigation [3,12].

After an inflammatory insult, including infection, the lung undergoes a complex process of tissue repair, which can result in deleterious fibrotic changes [13]. Fibrotic lung tissue loses much of its ability to regulate gas exchange between the air and the blood, frequently leading to organ failure. Interstitial lung disease (ILD) describes a broad range of typically inflammatory lung conditions characterized by lung fibrosis [14]. Due to the significant impact of fibrosis on pulmonary gas exchange, ILD is one of the leading indications for lung transplant in the United States [15]. γδ T cells may play a critical role in the progression of lung fibrosis. For example, after a bleomycin-induced lung injury in animal models, γδ T cells are enriched in the lung, incite tissue repair while delaying fibrotic changes in the lung [16–18]. However, little data exist regarding γδ T cell functionality in patients with ILD.

In several inflammatory conditions, murine γδ T cells are enriched in the lung and may migrate between tissue sites and their draining lymph node [19,20]. For example, we have previously reported that γδ T cells play a necessary role in regulating the neutrophil response in mouse models of severe pneumonia [21]. However, the functional role of γδ T cells in the human lung remains poorly understood. One recent report indicated broad phenotypic differences in human γδ T cells across a range of mucosal tissues, including the lung [22]. However, the functional relationship between γδ T cells in the human lung and its draining lymph nodes remains poorly understood, including how this relationship changes in inflammatory or fibrotic conditions. Within this context, we sought to investigate the phenotypic and functional differences between γδ T cells in the human lung and its draining hilar lymph nodes (HLN), leveraging matched tissue samples from patients with and without ILD. Understanding the functional differences between human γδ T cells in nonlymphoid tissues and lymph nodes, and how this functionality changes during inflammatory conditions, could provide critical knowledge into the role of γδ T cells within the respiratory mucosa.

## Materials and methods

### Tissue Collection and Processing

#### Deceased donor human samples

Potential deceased organ donors were identified nationally through the National Disease Research Interchange (NDRI). Patients without a history of lung disease, recreational substance inhalation, immunosuppression or total time of mechanical ventilation of more than 7 days were eligible for screening. Clinical and radiographic data of eligible donors were then reviewed by a board-certified physician for organ suitability including exclusion for evidence of current or recent infection, lung injury or warm ischemic time of > 60 minutes when applicable. Donors after either neurologic or cardiac death were accepted. At the time of procurement, whole lungs were recovered by the local Organ Procurement Organization in coordination with NDRI. Lungs were inflated to approximately 50% capacity with ambient air and the mainstem bronchus stapled. Lungs were then submerged into chilled histidine-tryphophan-ketoglutarate (HTK) fluid, secured and shipped to the University of Washington within 12-24 hours of procurement. Upon arrival, lung samples were inspected for evidence of gross pathology by a board-certified physician and inflated with ambient air prior to sample procurement. Lung samples of approximately 1 cm^3^ were then procured and immediately placed and stored in cold RPMI + 10% fetal bovine serum (FBS) media. Hilar lymph nodes were identified, dissected and stored in media. Samples were stored at 4°C until processing, which occurred within 24 hours.

#### ILD explant donor samples

ILD explant lung tissue was obtained in coordination with the University of Washington Lung Transplant Biorepository program. Informed consent was obtained from adults (age > 18 years) with a known diagnosis of ILD presenting for lung transplant. During lung transplant, the ILD lung was surgically removed by transplant surgeons and immediately delivered to the University of Washington Pathology Services. Lung and hilar lymph node samples were procured and stored in a similar fashion as noted above.

#### Ethical Approval

The processing of de-identified tissues from deceased donors was not considered to be human subjects research by the University of Washington IRB (STUDY00004036). Procurement and processing of tissues from patients with ILD undergoing lung transplant was approved under the University of Washington IRB (STUDY00008263).

#### Tissue single cell processing

Tissue single cell processing occurred within 1 calendar day of procurement. Tissue was minced into digestion media (RP7.5 [RPMI 1640 supplemented with 7.5% FBS, 2mM L-glutamine, 100 U/mL penicillin-streptomycin] + 700U/mL Collagenase Type II [Sigma Aldrich], 200U/mL DNase [Sigma Aldrich]) and placed in a shaking 37°C incubator for 50 minutes at 225 RPM. Cell suspensions were then washed with RP10 media (RPMI 1640 supplemented with 10% FBS, 2mM L-glutamine, 100 U/mL penicillin-streptomycin), strained through a 70µm filter, centrifuged at 400g for 5 minutes and washed again with RP10. Cells were treated with ACK lysis buffer to remove red blood cells, washed and strained again through a 70µm filter. For quantification, cell pellets were resuspended in RP10. For enumeration, single cell aliquots were analyzed on a Guava easyCyte (Cytek). Cells were then either immediately processed for flow analysis or cryopreserved in liquid nitrogen using cryopreservation solution (10% sterile DMSO (Sigma Aldrich), 90% FBS).

## Tissue Blocks for Immunohistochemistry

For histologic assessment, tissue samples were fixed in Formalin and then paraffin-embedded by the Fred Hutchinson Experimental Histopathology core. Paraffin-embedded blocks were sectioned and then stained. Slides from these blocks were stained with either H&E, Masson’s Trichrome or with fluorescent antibodies (Supplemental Table S1).

## Ex vivo stimulation

For stimulation experiments, 0.5–1×10^6^ cells per well were placed into 96-well V-bottom tissue culture plates. Cells were then cultured in RP10 media (RPMI 1640 supplemented with 10% FBS, 2mM L-glutamine, 100 U/mL penicillin-streptomycin). For stimulation, cells were cultured in RP10 containing either 1) phorbol myristate acetate (PMA, 50 ng/ml, Sigma) and ionomycin (500 ng/ml, Sigma) for 6 hours, 2) IL-12/15/18 (10 ng/mL) for 24 hours or 3) media alone (unstimulated). Cells were cultured at 37°C, 5% CO_2_ for 6 or 24 hours prior to flow cytometry processing. Four hours prior to sampling, GolgiPlug (BD Biosciences, 1:1000 dilution) and GolgiStop (BD Biosciences, 1:1500 dilution) was added for the remainder of the stimulation.

## Flow cytometric analysis

All flow panel reagent information, stain conditions, and gating are included in (Supplemental Figure S1, Supplemental Table S2). Of note, the spectral flow cytometry data for a subset of the deceased donor adult lung tissues (not including pediatric lungs, pediatric HLN or adult HLN tissues) were also included in a separate study and have been deposited on ImmPort (SDY3484) [23]. All flow staining was conducted at room temperature. For the viability staining, PBS was used for dilution. For FACS surface staining, FACSWash (1x PBS supplemented with 2% PBS) was used as the stain diluent. Surface stains were supplemented with Brilliant Stain Buffer Plus (BD Biosciences) for the 37-color spectral panel. For intracellular/nuclear staining, cells were initially fixed with the FOXP3 Fixation/Permeabilization Buffer Kit (Thermo Fisher) and then stained with antibodies using the FOXP3 Permeabilization Buffer (Thermo Fisher) as diluent. Cells were resuspended in FACSWash and events acquired on a FACSDiscover or a FACSSymphony. Flow cytometry data was analyzed using FlowJo v10 (BD Biosciences).

## Statistical analysis

Comparisons of matched samples was performed using either the paired t-test or Wilcoxon signed rank test. For unmatched comparisons, either t-test or Mann-Whitney test were used. P<0.05 was considered significant. All statistical testing was conducted using Prism v8 (GraphPad).

## Results

### γδ T cells are enriched in the lung in patients with and without inflammatory lung disease

To understand the lung-specific properties of γδ T cells, including during inflammatory conditions, we obtained matched lung and HLN tissues from both deceased donors without existing lung disease (N=11) and living donors with ILD undergoing lung transplant (N=10). Deceased donor (DD) tissues had a median age of 56 (interquartile range [IQR] 6-59; Supplemental Table S3). All deceased donors had a neurologic cause of death and a median duration of mechanical ventilation of 5 days (IQR 4-6). In the cohort of patients with ILD undergoing lung transplant, donors had a median age of 58 (IQR 52-65) and included a range of ILD-related diagnoses including idiopathic interstitial pneumonia (40%) and idiopathic pulmonary fibrosis (20%).

We first histologically assessed the lung tissue environments within a subset of the deceased donor (N=3) and ILD cohorts (N=3) by staining paraffin-embedded lung sections with H&E (**Figure 1A**) and a Masson’s Trichrome stain (**Figure 1B**). Lung and HLN samples from the deceased donor cohort showed relatively minimal inflammation, without substantial cellular infiltration and no evidence of excessive collagen deposition (**Figure 1A-B**). By contrast, lung samples from the ILD cohort demonstrated substantial lymphocytic infiltration, alveolar wall thickening and diffuse collagen deposition suggesting both acute and chronic inflammation and fibrotic changes (**Figure 1A-B**).

**Figure 1:**
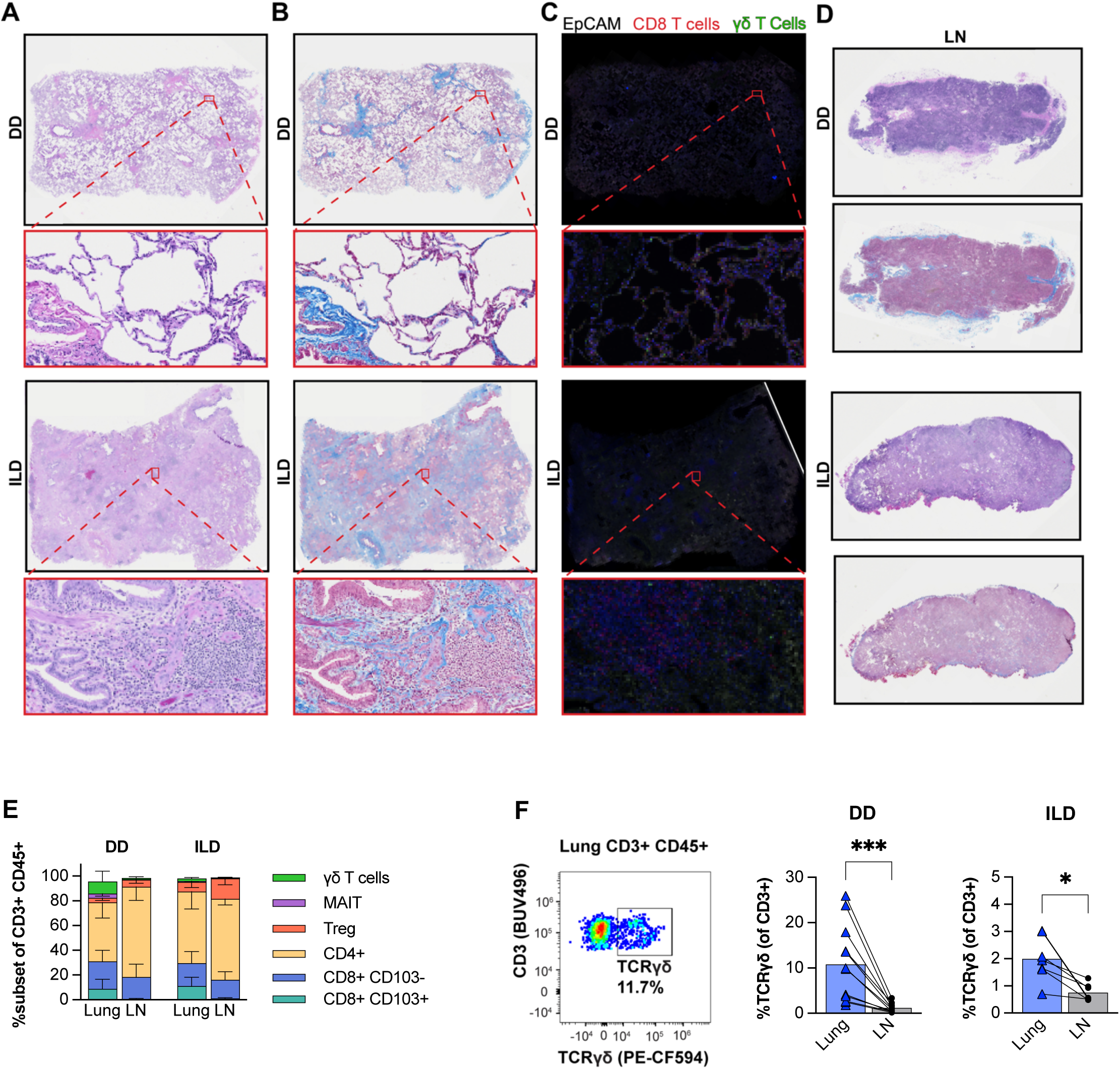
γδ T cells are enriched in the lung and retain spatial distribution during inflammatory lung disease. Lung and hilar lymph node (LN) tissues were obtained from deceased donors (DD) and patients with ILD undergoing lung transplantation. In 3 samples (both DD and ILD) lung and LN samples were stained with H&E (representative samples in A/D) or Masson’s Trichrome (representative samples in B). In addition, lung samples were stained with antibodies to EpCAM, CD8a and the γδ T cell receptor and examined by fluorescent microscopy (representative samples in C). Separately, fresh lung and LN samples were digested, stained with fluorescent antibodies and examined by flow cytometry. T cell subsets were identified (E) and the relative abundance of γδ T cells compared between matched lung and LN tissue (F). Comparisons by paired t-test; *P<0.05; ***P<0.001.

Given the substantial differences in lung tissue environment, we next sought to specifically identify the distribution of γδ T cells within these lung tissues using immunofluorescence. Lung sections were stained with antibodies for the γδ TCR, CD8 and epithelial cell adhesion molecule (EpCAM) and assessed by immunofluorescence (**Figure 1C**). γδ T cells were identifiable in both deceased donor as well as ILD lung tissue with similar spatial distribution within the interstitium. CD8^+^ cells displayed similar distribution patterns to γδ T cells in deceased donor lung tissue but appeared in clusters in ILD lung tissue.

To further assess the relative abundance of γδ T cells and other T cell subsets in the lung we next used cryopreserved single cell suspensions from lung and matched HLN for analysis by flow cytometry. γδ T cells were identified after gating for CD3^+^ γδ TCR^+^ cells within the CD45^+^ cell compartment (Supplemental Figure S1A). Flow cytometry samples that yielded less than 30 γδ T cell events were not included in phenotypic and functional analyses. In line with the histologic assessments, γδ T cells were identified in both lung and HLN tissue, regardless of disease state (**Figure 1 E-F**). Notably, γδ T cells were more abundant within the T cell population of the lung compared to HLN in both the deceased donor (7.2% lung vs 0.6% HLN, P=0.002) and ILD tissues (2.0% lung vs 0.8% HLN, P=0.016). Given differences in patient population as well as tissue procurement and processing, we did not directly compare γδ T cell frequency between deceased donor and ILD lung tissues, though we observed a trend towards decreased frequency of γδ T cells in the T cell compartment in ILD lung tissue compared to deceased donors.

A previous study reported the frequency of γδ T cells within the T cell compartment of the lung is nearly twice as high in pediatric compared to adult lungs [22]. We observed a similar relative γδ T cell frequency in children (< 10 years) compared to adults (>18 years) (Supplemental Figure S2A). While γδ T cells were more abundant in the T cell compartment in pediatric (N=5) compared to adult lungs (N=8, P<0.001), the relative abundance of γδ T cells in the lungs of both groups were higher compared to matched HLN tissue (P<0.05, both; Supplemental Figure S2A). Taken together, these data suggest γδ T cells are enriched in the T cell compartment of the lung compared to regional lymph nodes, regardless of inflammatory state.

### γδ T cell phenotype differs between lung and regional lymph nodes

We next investigated the impact of tissue type and inflammatory state on γδ T cell phenotype using flow cytometry. To control for potential batch effects, a technical control (standardized aliquots from a single leukapheresis donor) was included in all experiments to allow for direct comparisons within cohorts over time. Additionally, given the inherent differences in procurement processes between the deceased donor and ILD cohorts, we continued to employ a conservative analysis approach by specifically comparing relative differences in cell populations in the lung compared to matched HLN tissue.

We first assessed CCR7 and CD45RA expression to identify populations with a naïve (CCR7^+^ CD45RA^+^), effector memory (Tem: CCR7^-^ CD45RA^-^), central memory (Tcm: CCR7^+^ CD45RA^-^) and CD45RA-expressing effector memory (CCR7^-^ CD45RA^+^; these have also been referred to as recently activated effector memory cells and terminally differentiated effector memory cells: TEMRA) phenotype. We found the majority of γδ T cells expressed an effector memory (Tem or TEMRA) phenotype, regardless of tissue type or presence of ILD **(Figure 2A)**. We observed a population of γδ T cells with a naïve phenotype in HLN, while the γδ T cell population in deceased donor and ILD lung consisted nearly exclusively of TEMRA and Tem cells. Notably, the frequency of TEMRA γδ T cells was higher in deceased donor lung tissue compared to HLN (77% lung vs 52% HLN, P<0.001), a difference consistent across age ranges (Supplemental Figure S2B), and not observed in ILD tissues (60% lung vs 48% HLN, P=0.09, **Figure 2A**).

**Figure 2:**
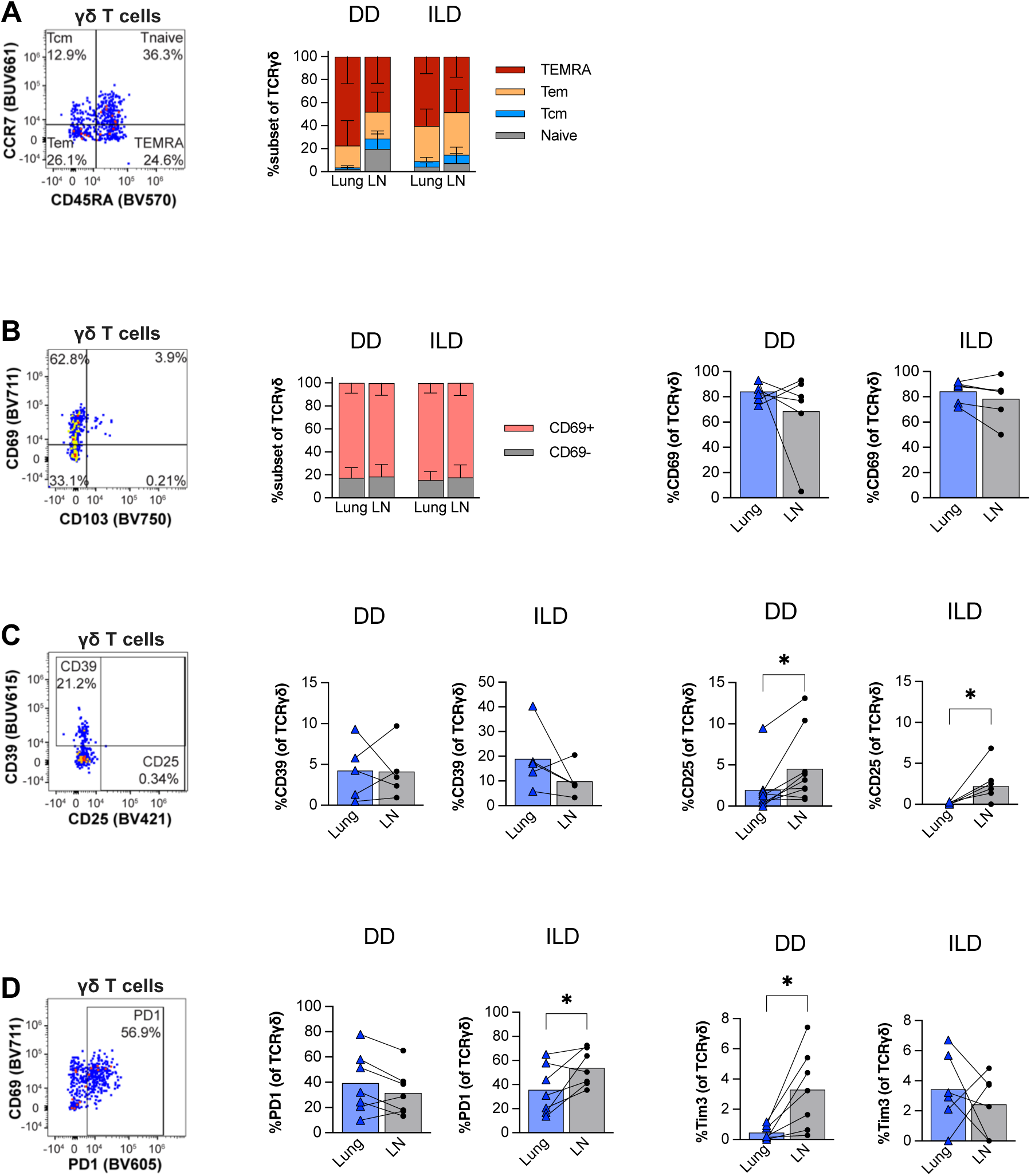
Lung γδ T cells demonstrate tissue-specific effector memory and activation phenotypes. Lung and hilar lymph node (LN) tissues were obtained from deceased donors (DD) and patients with ILD undergoing lung transplantation. Matched lung and LN samples were digested, stained with fluorescent antibodies and examined by flow cytometry. γδ T cell phenotypes including memory subsets (A) as well as expression of CD69 (B), CD39 and CD25 (C) and PD1 and Tim3 (D). Comparisons of matched samples was performed using either the paired t-test or Wilcoxon signed rank test; *P<0.05.

We next investigated biomarkers that could indicate γδ T cell tissue-residency in both lung and HLN given the possibility of cell migration between lymph and mucosal sites [23]. γδ T cells in the lung and HLN had similar expression patterns of CD69, regardless of the presence of inflammatory lung disease (**Figure 2B**). While ∼80% of γδ T cells expressed CD69 regardless of tissue location or disease state, only a small fraction expressed CD103.

We next assessed activation markers including CD25 (IL-2Ra chain) and CD39 (an ectonucleotidase, involved in the conversion of ATP to adenosine). CD39 expression by γδ T cells was relatively conserved regardless of tissue type or inflammatory state (**Figure 2C**). Conversely, γδ T cell expression of CD25 was overall low in deceased donor lungs and undetectable in ILD lungs, though increased in HLN compared to lung tissues in both deceased donors (P<0.01) and ILD patients (P<0.05). Given these findings of decreased relative activation of γδ T cells in ILD tissues, we next assessed PD-1 expression, a biomarker often associated with functional exhaustion but also induced by TCR stimulation and pro-inflammatory cytokines [24]. PD-1 expression did not differ between lung and HLN γδ T cells in either the deceased donor or ILD cohorts (**Figure 2D**). However, we observed a trend towards increased PD-1 expression in ILD HLN γδ T cells (P=0.05) relative to matched lung resident γδ T cells, potentially suggesting the presence of activating stimuli. Conversely, expression of Tim3, another biomarker of exhaustion, was detected on a small population of γδ T cells, and significantly higher in HLN vs. lung γδ T cells in deceased donors (P=0.02) but not ILD patients. Overall, these phenotypic data indicate that γδ T cells within the lung tissue have distinct phenotypes compared to regional lymph nodes, regardless of inflammatory conditions.

### γδ T cell functional properties differ between lung and regional lymph nodes

Our phenotypic assessment of γδ T cells suggested differences based on tissue type and inflammatory state. Therefore, we next investigated whether tissue type and inflammatory state affect γδ T cell functional properties. Single cell suspensions of lung and HLN tissue were stimulated for 6 hours using PMA/ionomycin and compared to unstimulated samples (**Figure 3**). Stimulated γδ T cells in humans can produce a variety of cytokines, including IFNγ, TNFα and IL-17 [21,22]. We identified relatively low IL-17 expression by γδ T cells, regardless of tissue type or inflammatory state (Supplemental Figure S3). Conversely, stimulated γδ T cells demonstrated increased expression of both IFNγ and TNFα in lung and HLN deceased donor tissues (**Figure 3A**). However, in ILD tissues, HLN γδ T cells retained higher capacity to express both IFNγ and TNFα (both P>0.01) compared to γδ T cells from ILD lung tissue. No difference was observed in IFNγ and TNFα expression by effector memory state, regardless of tissue type or inflammatory state.

**Figure 3:**
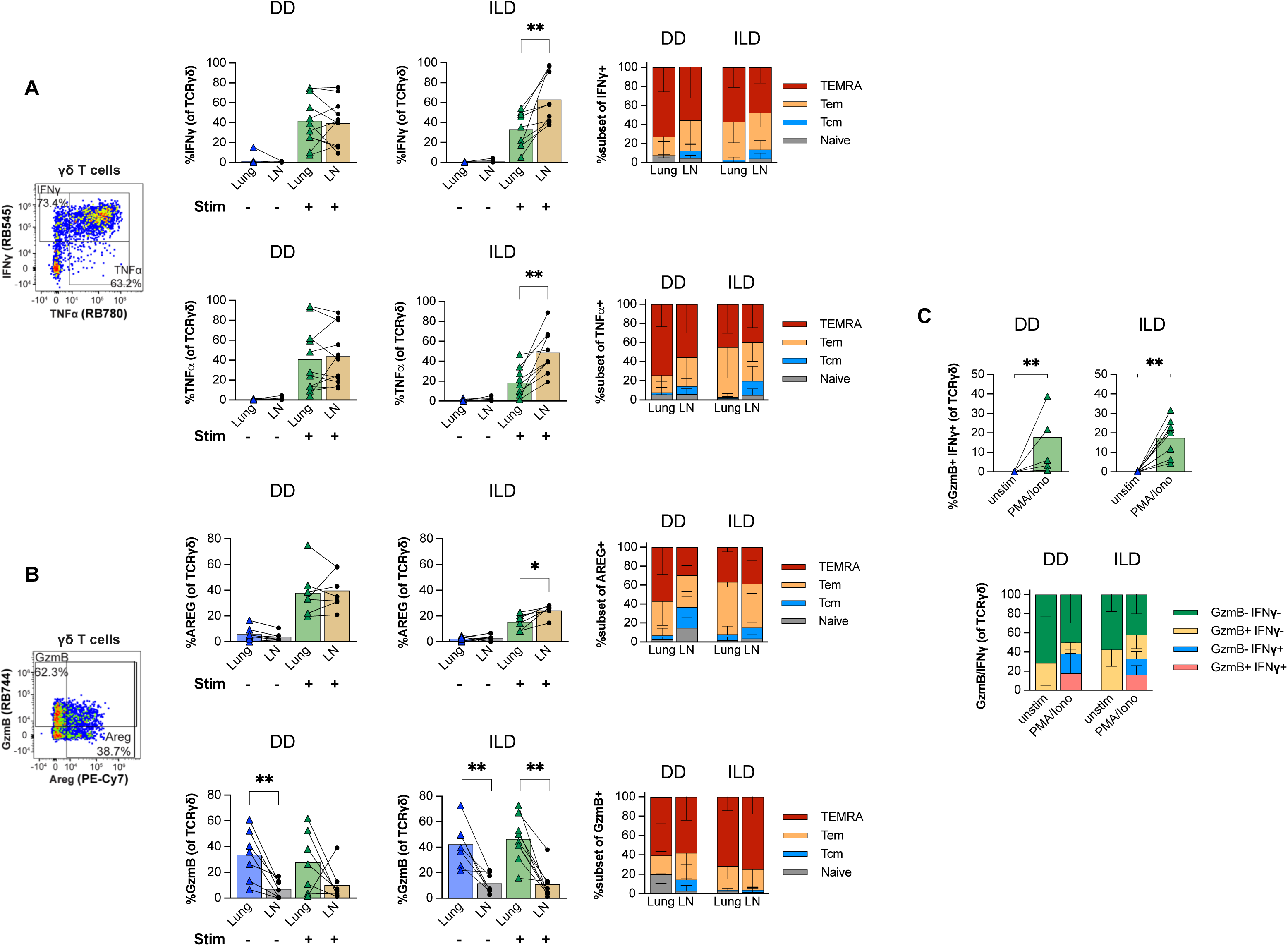
Lung γδ T cells express increased cytolytic capacity relative to regional lymph nodes. Lung and hilar lymph node (LN) tissues were obtained from deceased donors (DD) and patients with ILD undergoing lung transplantation. Matched lung and LN samples were digested, stimulated with PMA/ionomycin for 6 hours, stained with fluorescent antibodies and examined by flow cytometry. Differences in intracellular IFNγ or TNFα (A) as well as GzmB or AREG (B) as well as relative expression in memory subsets were compared between matched tissues before and after stimulation. Lung γδ T cell expression of both GzmB and IFNγ was compared before and after PMA/ionomycin stimulation (C). Comparisons of matched samples was performed using either the paired t-test or Wilcoxon signed rank test; *P<0.05. **P<0.01.

γδ T cells can express amphiregulin (AREG), a protein associated with tissue repair as well as pulmonary fibrosis [25–27]. Given the potential relevance of AREG in ILD, we assessed expression before and after PMA/ionomycin stimulation. Without stimulation, γδ T cells showed minimal AREG expression (**Figure 3B**). However, after stimulation, γδ T cells had similar AREG expression in deceased donor lung (38%) and HLN tissues (40%), with the majority of AREG-expressing cells having an effector memory or TEMRA phenotype. Notably, we observed not only a trend towards overall lower γδ T cell AREG expression in ILD tissues but relatively more frequent AREG expression in HLN (24%, P=0.03) compared to lung (16%) γδ T cells. The majority of AREG-producing γδ T cells had either a TEMRA or effector memory phenotype.

Human γδ T cells can express and release granzyme B (GzmB), with cytolytic capacity described in animal models [22,28]. We observed that PMA/ionomycin stimulation for 6 hours was not sufficient to increase γδ T cell GzmB expression (**Figure 3B**). However, in unstimulated cells, lung γδ T cells from both deceased donors (34%) and ILD patients (37%) had relatively higher expression of GzmB compared to those in the HLN (deceased donors: 7%; ILD: 10%; P=0.01, both, **Figure 3B**), which is in line with the increase in effector memory cells in the lungs and may further indicate tissue residency. We did not observe a difference between pediatric and adult γδ T cell GzmB expression (Supplemental Figure S3D). Finally, we assessed the co-expression of both GzmB and IFNγ capacity after stimulation. We observed a comparable increase in GzmB^+^ IFNγ^+^ γδ T cells after PMA/ionomycin stimulation (**Figure 3C**). Overall, our ex vivo stimulation data indicate that the γδ T cell population has a comparable functional capacity in deceased donor lungs and HLN, while γδ T cells from ILD lungs displayed a decreased functional capacity compared to their HLN counterparts

### Ex vivo cytokine stimulation yields polyfunctional responses in lung γδ T cells regardless of inflammatory state

We wanted to further interrogate the potential functional impairment of γδ T cells in ILD lungs. We reasoned that a stimulation signal that is less potent than PMA/ionomycin may reveal potential functional deficits. We thus stimulated single cell suspensions from deceased donor and ILD lungs for 24 hours with either media or a cytokine combination of IL-12, IL-15 and IL-18 (IL-12/15/18, **Figure 4A**). Matched HLN cells were not available to include in these cytokine stimulation experiments. γδ T cells from both deceased donor and ILD lungs demonstrated increased expression of GzmB after IL-12/15/18 stimulation (**Figure 4B**). Additionally, IL-12/15/18 stimulation resulted in increased expression of IFNγ, though not TNFα or AREG, in γδ T cells from both deceased donor and ILD lungs (**Figure 4B**). These results indicate inflammatory cytokines specifically stimulated GzmB and IFNγ in lung γδ T cells, a functionality retained in ILD. We next assessed the relative expression of both GzmB and IFNγ capacity after the 24-hour IL-12/15/18 stimulation. The majority of IFNγ-producing γδ T cells also expressed GzmB after cytokine stimulation, regardless of donor cohort (**Figure 4C**). Thus, the cytokine stimulation did not reveal any functional defects, largely mirroring the PMA/ionomycin findings [29,30]. Finally, we sought to understand whether polyfunctional responses, including simultaneous cytokine expression, were present in cells co-expressing GzmB and IFNγ. Although the cytokine stimulation produced relatively limited TNFα expression in the broader population γδ T cells, we observed TNFα expression in the GzmB^+^ IFNγ^+^ γδ T cells in both the deceased donor and ILD lungs (**Figure 4D**). As we further explore in the discussion, lymphocytes capable of simultaneous effector molecule expression may have particular importance during inflammatory conditions or at mucosal barriers such as the lung [29,30]. Overall, our findings suggest that γδ T cells in lung and LN are capable of polyfunctional responses that are conserved even during chronic inflammatory lung conditions.

**Figure 4:**
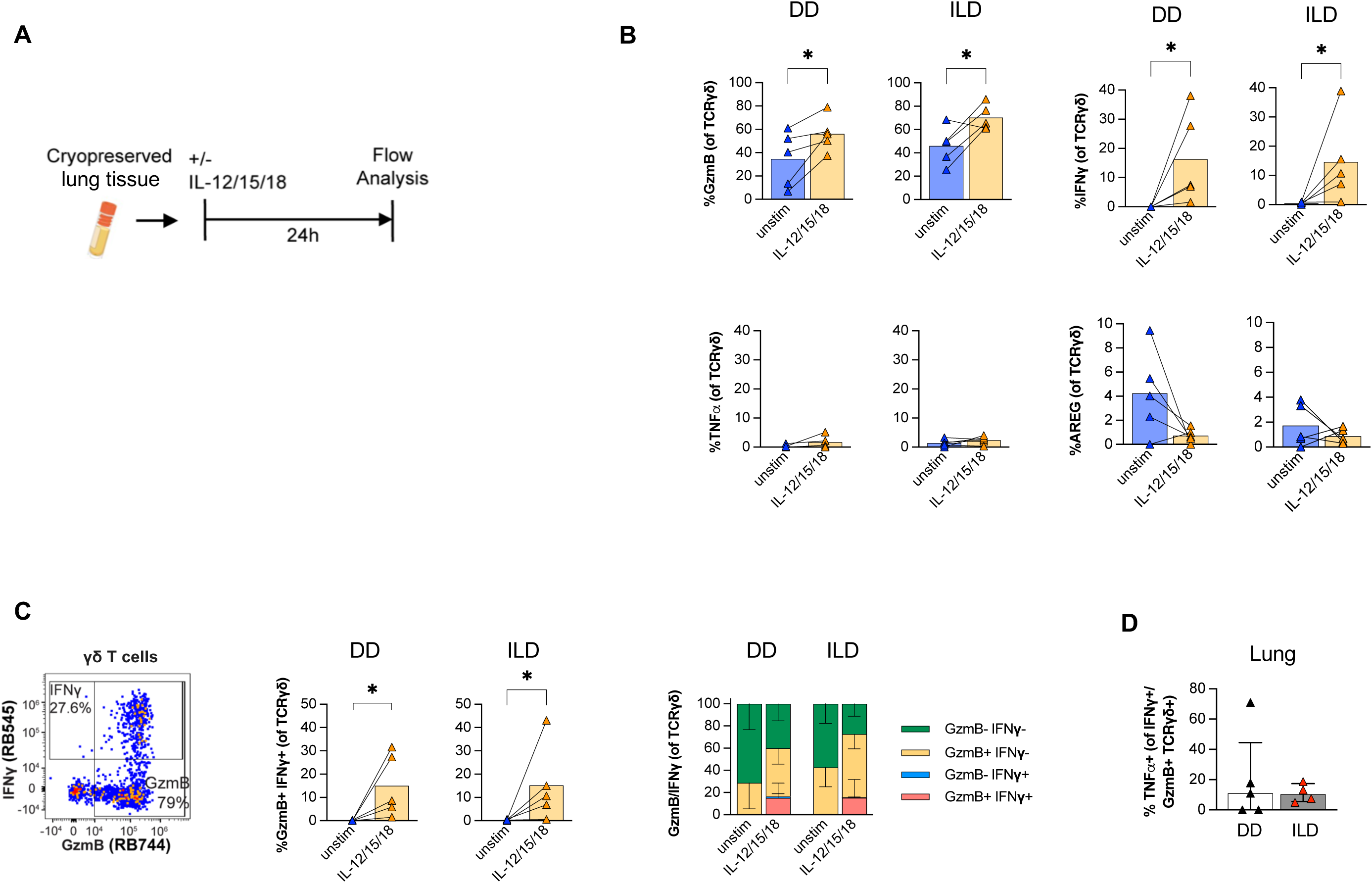
Functional capacity is retained in lung γδ T cells during chronic inflammatory lung disease. Lung tissues were obtained from deceased donors (DD) and patients with ILD undergoing lung transplantation. Tissue samples were digested and cryopreserved. Cells were later stimulated with IL-12/15/18 for 24 hours, stained with fluorescent antibodies and examined by flow cytometry (A). Lung γδ T cell expression of GzmB, IFNγ, TNFα and AREG were compared before and after stimulation (B). Lung γδ T cell expression of both GzmB and IFNγ was compared before and after 24-hour IL-12/15/18 (C). Expression of TNFα in GzmB^+^ IFNγ^+^ lung γδ T cell after 24-hour IL-12/15/18 (D). Samples yielding less than 25 cell events were not included in Figure D. Comparisons between unstimulated and stimulated cells were made using either the paired t-test or Wilcoxon signed rank test; *P<0.05. **P<0.01.

## Discussion

The lung, with a complex architecture and microbiome, serves as a unique mucosal environment where tissue inflammation can result in fibrotic lung disease. γδ T cells are abundant in the mucosa early after development, with increasingly described roles in immune responses to both commensal and pathogenic organisms [4,31]. However, human γδ T cells demonstrate profound phenotypic variability between individuals and tissues, and little is known regarding the phenotype and function of γδ T cells within the human lung, regardless of disease state [21,22]. Given this knowledge gap, we compared matched lung and HLN tissue from both previously healthy deceased donors as well as living donors with ILD. We then determined whether the phenotypic and functional capacities of human γδ T cells are tissue and disease-dependent. We found that γδ T cells are enriched within the T cell compartment in the lung and demonstrate unique functionality compared to its draining lymph node. Intriguingly, we also found that much of the lung γδ T cell functionality, including cytotoxic and cytokine properties, is conserved, even during severe fibrotic lung disease.

Within the mucosal microenvironment, γδ T cells may regulate tissue inflammation including propagation, de-escalation as well as subsequent tissue repair [6,7]. However, the functional role of γδ T cells within the lung has been largely confined to animal models, with limited data in humans. Furthermore, human γδ T cells differ substantially from their murine counterparts, including in TCR subsets as well as functionality [32]. γδ T cells are predominantly composed of the Vδ1 subset in both adult human lungs and its associated lymph nodes [22]. We found that γδ T cells are relatively more abundant in the lung mucosa and acquire a specific cytolytic capacity compared to their regional lymph node counterparts-as evidenced by profoundly upregulated GzmB expression. This difference in GzmB expression could be partially related to the proportional increase of effector memory phenotype γδ T cells in the lung or could be an indicator of tissue residence, particularly in context of the associated CD69 expression. The finding that such a large fraction of γδ T cells in the lung express GzmB could also reflect a critical cytotoxic role of γδ T cells within the mucosal barrier of the lung [33,34]. γδ T cells can produce several cytotoxic products, including granzymes, perforin and granulysin, and have been investigated for their role in cancer pathogenesis and potential for therapeutics [35]. The ability to lyse infected cells may also serve as a necessary role for γδ T cells within the mucosal immune system, and γδ T cell-mediated cell lysis has been implicated in host defense against intracellular viral, parasitic and bacterial infections [36–38].

In mice, γδ T cells, including in the lung, have been described to produce IFNγ, TNFα and IL-17 in certain inflammatory conditions [21,39]. However, our findings suggest stimulation of both human lung and HLN γδ T cells results in limited IL-17 expression. Conversely, we report lung γδ T cells produce IFNγ after either broad PMA/ionomycin or IL-12/15/18 cytokine stimulation. Indeed, inflammatory stimulation of lung γδ T cells with IL-12/15/18 primarily increased IFNγ production in cells also expressing GzmB. Notably, this polyfunctional capacity was apparent after IL-12/15/18 stimulation, in which a population of GzmB^+^ IFNγ^+^ TNFα^+^ γδ T cells were observable in the lung and retained even in severely diseased lung with ILD. Lymphocyte polyfunctionality, the ability to express multiple effector proteins simultaneously, has become increasingly recognized as a critical trait during inflammatory conditions, including infection and cancer [40–43]. Limited data exist regarding γδ T cell polyfunctionality, though GzmB^+^ IFNγ^+^ TNFα^+^ populations have been reported in circulating and mucosal γδ T cell subsets in humans [22]. Suggesting a potential role in immune responses to infection, an expanded population of GzmB^+^ IFNγ^+^ γδ T cells has also been reported in patients with tuberculosis [44]. Therefore, the conservation of both cytolytic (GzmB) and cytokine (IFNγ/TNFα) capacity in lung γδ T cells even during chronic inflammation could be represent a protective role against mucosal infection. For example, γδ T cells produce IFNγ in response to multiple viral and intracellular bacterial stimuli [45–47]. Similarly, GzmB-mediated cytotoxicity has been reported by γδ T cells in models of viral and bacterial pathogens [48,49]. Indeed, we observed that γδ T cell expression of both IFNγ and TNFα in patients with ILD differs compared to matched draining lymph nodes. While this finding may represent increased responsiveness in lymph node γδ T cells during inflammation, it may also represent impaired responses in the lung. Therefore, the maintenance of polyfunctionality may be crucial during periods of prolonged lung inflammation, when mucosal barriers may be disrupted.

An intriguing finding from this study concerns the decreased ability of the lung γδ T cell population to express AREG in patients with ILD compared to healthy controls. γδ T cell-derived AREG-an epidermal growth factor involved in fibroblast proliferation-may be important in mucosal tissue repair and barrier integrity, including in murine lung infection [25,50]. AREG may also mediate lung fibrosis, with circulating levels increased in patients with ILD and excessive alveolar epithelial cell AREG-production potentially contributing to ILD pathogenesis [26,27]. The role of γδ T cells in ILD has not been well deciphered. For example, after a bleomycin-induced lung injury in mouse models, γδ T cells are enriched in the lung and incite tissue repair while delaying fibrotic changes in the lung [16–18,51]. Conversely, IL-17-producing cells, such as murine γδ T cells, may also activate fibroblasts, propagating fibrosis [52]. However, our finding that γδ T cell AREG-capacity is maintained in healthy lungs but reduced in end-stage ILD may suggest a role of γδ T cells in the homeostasis of the lung mucosa. As all ILD lungs were procured at the time of lung transplantation, implying chronic disease, it’s also possible different γδ T cell phenotypes may be observed earlier in the disease course, during periods of acute inflammation or after exposure to immunomodulating agents.

The effects of a chronic inflammatory lung environment on lymphocyte function are not well understood. However, we observed that in healthy controls, lung and HLN γδ T cells expressed similar functional capacity including stimulated expression of IFNγ, TNFα and AREG. Conversely, lung γδ T cells from ILD patients consistently had lower stimulated expression of IFNγ, TNFα and AREG relative to matched HLN after PMA/ionomycin stimulation, However, lung γδ T cells retained a robust cytokine response after IL-12/15/18 stimulation in both healthy controls and ILD tissues, which could be due to the inherently different nature of these activating signals as well as the difference in stimulation duration. These findings indicate relatively preserved functional capacity, even during chronic inflammatory lung disease.

Our study has several strengths. We are the first study, that we are aware, to functionally characterize γδ T cells in healthy and diseased human lung relative to matched regional lymph node tissues. This approach, allowing for subject-specific internal comparisons, accounts for variations in subject characteristics, procurement and processing. Furthermore, all included subjects underwent extensive screening by a physician prior to tissue procurement and subsequent processing. Additionally, we utilized a highly optimized flow cytometry approach as well as accounted for batch-to-batch variability. A limitation of our study is that we do not directly assess the differences in TCR-based Vδ1 and Vδ2 subset phenotype and function between tissues. However, the Vδ1 subset has previously been reported to predominate with similar proportions in both adult human lung and lung-associated lymph node tissue, and TCR-based γδ T cell subsets do not fully reflect functional differences [22]. Therefore, to allow for a more complete assessment of γδ T cells within the lung and its regional lymph nodes, we identified all γδ T cells and then compared phenotypic and functional changes based on tissue type and inflammation.

Overall, we report that lung-specific polyfunctionality is retained even during a severe fibrotic disease state. Our data also provide evidence regarding the evolution of mucosa-relevant characteristics as γδ T cells migrate between the lung and regional lymph system. Furthermore, the conservation of certain phenotypes during pulmonary fibrosis necessitates further study to elucidate the role of γδ T cells in the pathogenesis of chronic, inflammatory lung diseases.

## CRediT authorship contribution statement

**Alexis Taber**: Writing – review & editing, Writing – original draft, Methodology, Investigation, Formal analysis, Data curation, Visualization. **Marie Frutoso**: Writing – review & editing, Methodology, Investigation, Formal analysis, Data curation. **Nicole Potchen**: Writing – review & editing, Formal analysis, Data curation. **Amanda Koehne**: Writing – review & editing, Methodology, Investigation. **Chelsea Schmitz**: Writing – review & editing, Resources, Investigation. **Eric D. Morrell**: Writing – review & editing, Resources, Funding acquisition. **Martin Prlic**: Conceptualization, Supervision, Project administration, Funding acquisition, Writing – review & editing, Writing – original draft, Methodology, Formal analysis, Visualization, Data curation. **Shelton W. Wright**: Conceptualization, Supervision, Project administration, Funding acquisition, Writing – review & editing, Writing – original draft, Methodology, Visualization, Investigations, Resources, Formal analysis, Data curation.

## Acknowledgements

We would like to thank members of the Prlic and Wright labs for critical discussion. We thank Andrew Konecny, Eva Domenjo, Guilhem Rerolle and Abdullah Bashmail for technical assistance with experiments. We also thank the Fred Hutch Experimental Histopathology core and Flow Cytometry core. We also thank T. Eoin West at the University of Washington for access to deceased donor tissues. Finally, we would like to thank the patients and families who donated their organs for this work, and we acknowledge the use of tissues procured by the National Disease Research Interchange (NDRI; RRID:SCR_000550). This work was supported by the US National Institutes of Health [grants R01AI123323 (MP), K12HD047349 (SWW), K08HL157562 (SWW), R21AI130797 (to T. Eoin West for tissue procurement) and R01HL169265 (EDM)]. Additionally, this work was supported by NIH P30 CA015704 of the Fred Hutch/University of Washington/Seattle Children’s Cancer Consortium, which includes the Experimental Histopathology Shared Resource, RRID:SCR_022612, and the Flow Cytometry Shared Resource, RRID:SCR_022613. Finally, the Fred Hutch Experimental Histopathology Shared Resource equipment is supported by a grant from the M.J. Murdock Charitable Trust grant SR-202221337.

