## Supplemental Figures for "Human lung γδ T cells maintain functionality during inflammatory lung disease"

### Supplementary Data

**Supplementary Figure S1. Gating strategy.** Representative gating strategy for γδ T cells, **A)** Spectral panel and **B)** Symphony Panel.

**A** Spectral Panel:

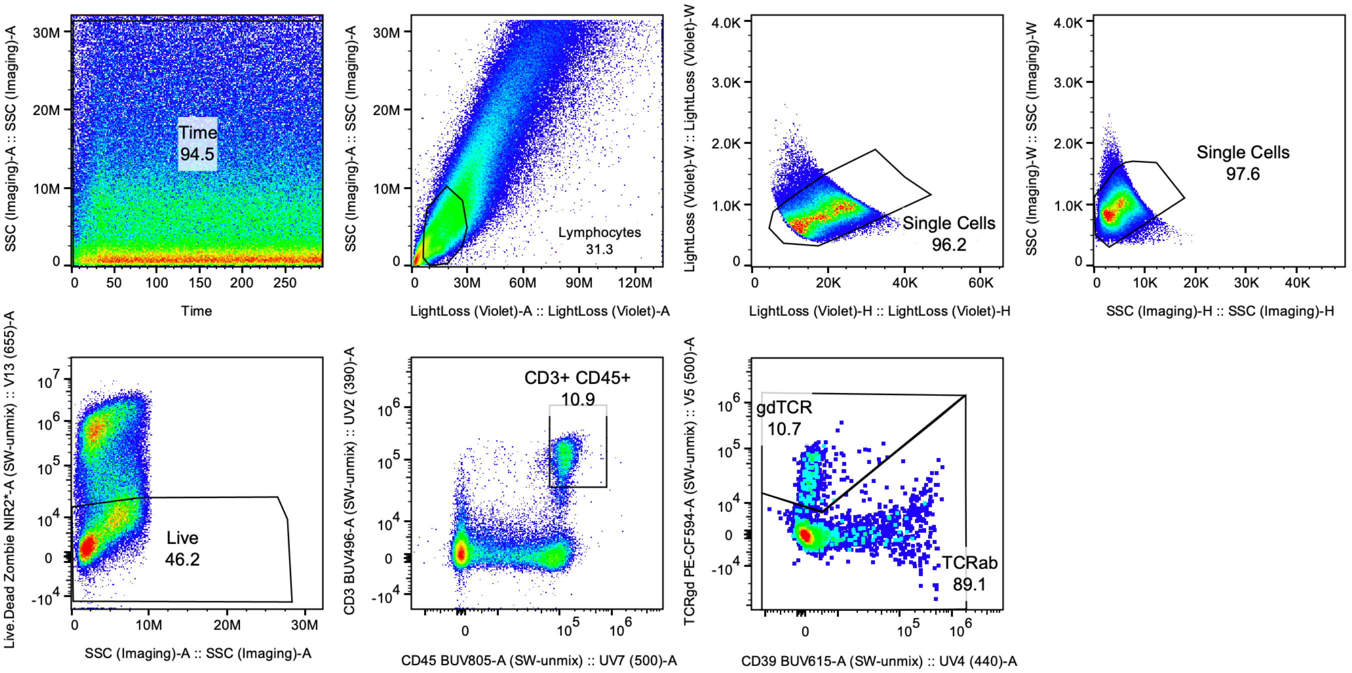


**B** Symphony Panel:

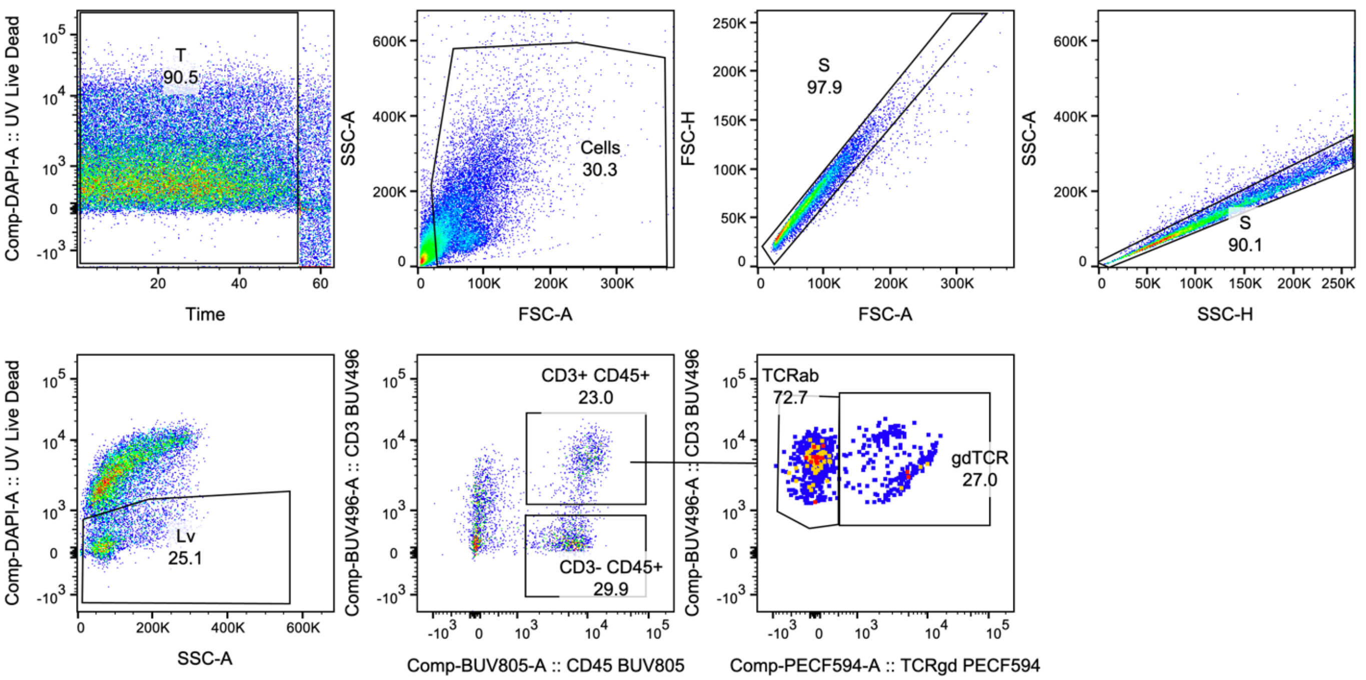

**Supplementary Figure S2. Comparison of deceased donor lungs by age.** Unstimulated pediatric (< 10 years of age) and adult (>18 years of age) lung and hilar lymph node samples from deceased donors samples gated for **A)** γδ T cells, **B)** γδ T cell memory phenotypes, **C)** CD69 and **D)** granzyme B. Samples were stimulated with PMA/ionomycin for 6 hours and stained for **E)** γδ T cell intracellular IFNγ and TNF. Comparisons made by paired and unpaired t-test.

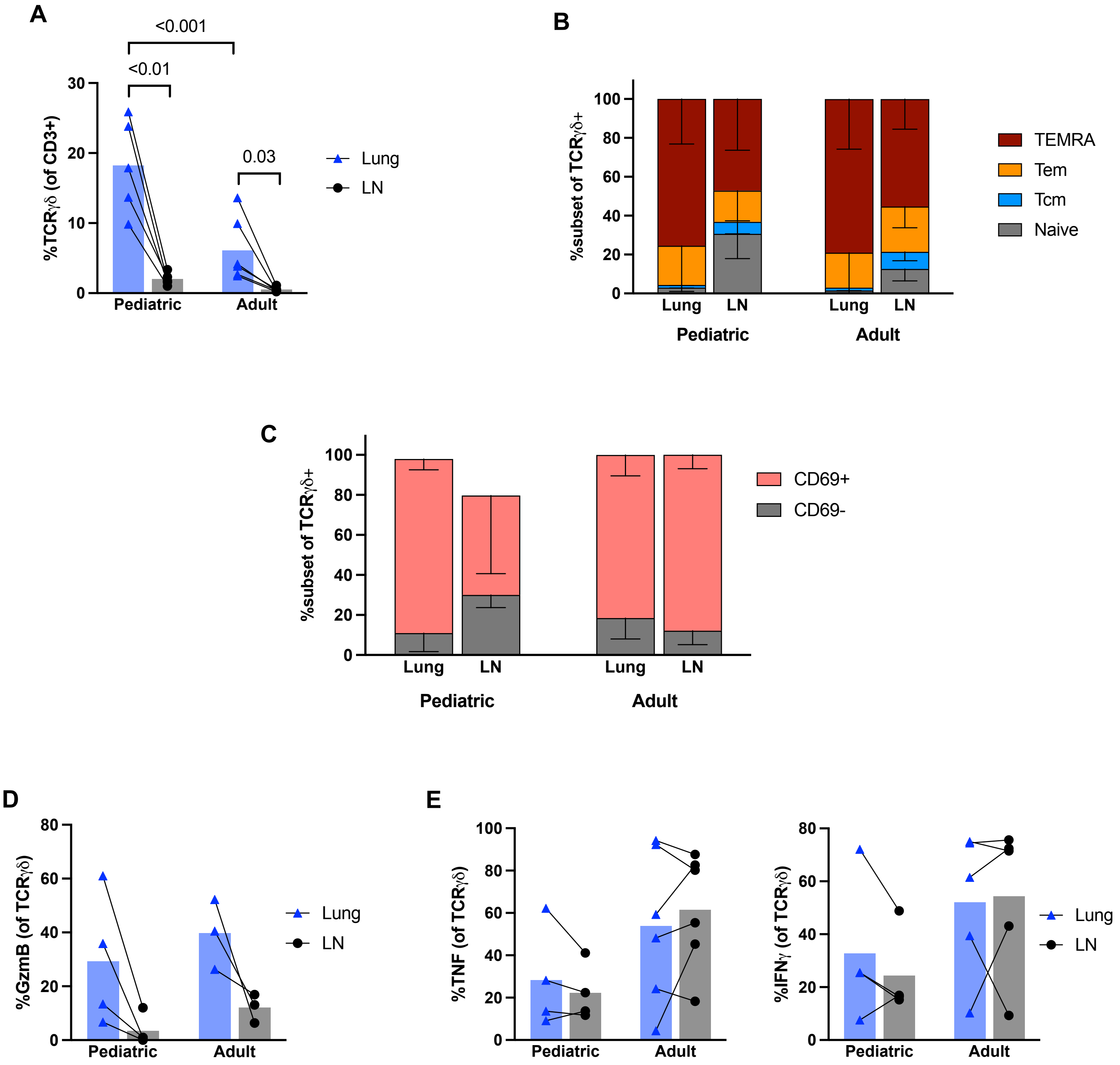

**Supplemental Figure S3. Phenotype and cytokine expression by γδ T cells.** Single cell suspensions were stimulated with PMA/Ionomycin or media alone for 6 hours, stained with antibodies and percentage of specific marker expression assessed by flow cytometry after gating for γδ T cells. **A**) Unstimulated and PMA/ionomycin stimulated cells with specific marker expression. **B**) Heatmap displaying the mean expression of unstimulated γδ T cells for specific markers. DD: deceased donor; ILD: interstitial lung disease donor; LN: hilar lymph node; stim: PMA/ionomycin stimulation. Statistical analysis by paired t-test or Wilcoxon test. *P<0.05, **P<0.01.

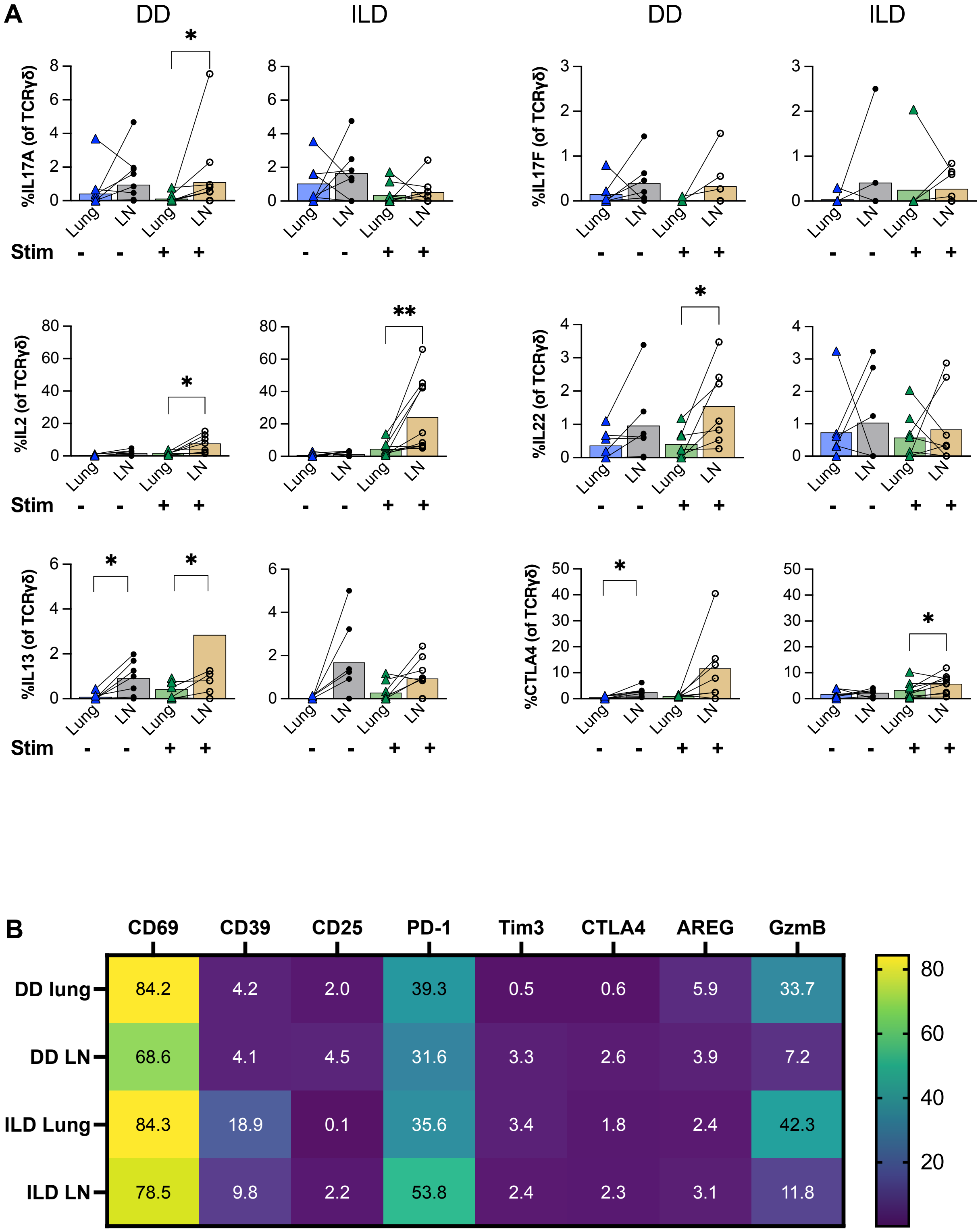

**Supplementary Table S1: Immunohistochemistry reagent panel**

| **Lungs** | | | | | |
|---|---|---|---|---|---|
| **Antibody** | **Clone** | **Host** | **Manufacturer Catalog #** | **Concentration** | **Fluor** |
| EpCAM | Ber-Ep4 | Mouse | Biolegend 324202 | 0.5 ug/mL | 570 |
| gdTCR | H-41 | Mouse | Santa Cruz sc-100289 | 0.5 ug/mL | 520 |
| CD8a | 144B | Mouse | Dako M0755 | 0.2 ug/mL | 690 |
| **Lymph Nodes** | | | | | |
| **Antibody** | **Clone** | **Host** | **Manufacturer Catalog #** | **Concentration** | **Fluor** |
| CD20 | L-26 | Mouse | Dako M0755 | 12 ug/mL | 570 |
| gdTCR | H-41 | Mouse | Santa Cruz sc-100289 | 0.5 ug/mL | 520 |
| Podoplanin | Polyclonal | Rabbit | Sigma HPA007534 | 0.1 ug/mL | 690 |

### Supplementary Table S2: Flow Cytometry panels

#### 37 color Spectral T cell panel

| | S8 Peak Detector | Fluorophore | Antigen | Cocktail | Dilutions | |
|---|---|---|---|---|---|---|
| 1 | UV01 [375] (0.98), UV03 [420] (1.00) | Spark UV 387 | CD4 | *Intracellular* | 1: | 80 |
| 2 | UV03 [420] (1.00) | BUV395 | CD8a | Surface | 1: | 80 |
| 3 | UV06 [460] (1.00) | Autofluorescence | - | NA | 1: | NA |
| 4 | UV07 [500] (1.00) | BUV496 | CD3 | *Intracellular* | 1: | 40 |
| 5 | UV11 [575] (1.00) | BUV563 | CD161 | Surface | 1: | 20 |
| 6 | UV13 [605] (0.85), YG03 [605] (1.00) | BUV615 | CD39 | Surface | 1: | 20 |
| 7 | UV15 [655] (0.89), R1 [655] (1.00) | BUV661 | CCR7 | Surface | 1: | 10 |
| 8 | UV18 [725] (1.00) | BUV737 | ICOS | Surface | 1: | 160 |
| 9 | UV21 [810] (0.65), UV03 [420] (1.00) | BUV805 | CD45 | Surface | 1: | 80 |
| 10 | V01 [420] (1.00) | BV421 | CD25 | Surface | 1: | 40 |
| 11 | V03 [460] (1.00) | V450 | IL-17F | *Intracellular* | 1: | 20 |
| 12 | V04 [475] (1.00) | BV480 | CD28 | Surface | 1: | 40 |
| 13 | V06 [515] (1.00) | BV510 | CD27 | Surface | 1: | 20 |
| 14 | V09 [575] (1.00) | BV570 | CD45RA | Surface | 1: | 160 |
| 15 | V11 [605] (1.00) | BV605 | PD1 | Surface | 1: | 20 |
| 16 | V13 [655] (1.00) | BV650 | Ki67 | *Intracellular* | 1: | 320 |
| 17 | V15 [700] (1.00) | BV711 | CD69 | Surface | 1: | 320 |
| 18 | V17 [750] (1.00) | BV750 | CD103 | Surface | 1: | 160 |
| 19 | V18 [780] (1.00) | BV785 | CD127 | Surface | 1: | 10 |
| 20 | B02 [515] (1.00) | BB515 | TIM3 | Surface | 1: | 80 |
| 21 | B04 [545] (1.00) | RB545 | IFNg | *Intracellular* | 1: | 20 |
| 22 | B07 [605] (1.00) | BB630-P2 | CTLA4 | *Intracellular* | 1: | 80 |
| 23 | B09 [655] (0.53), R01 [655] (1.00) | BB660-P2 | IL-13 | *Intracellular* | 1: | 640 |
| 24 | B11 [700] (0.90), R03 [700] (1.00) | BB700 | IL-22 | *Intracellular* | 1: | 640 |
| 25 | B13 [750] (1.00) | RB744 | Granzyme B | *Intracellular* | 1: | 320 |
| 26 | B14 [780] (1.00) | RB780 | TNFa | *Intracellular* | 1: | 40 |
| 27 | YG01 [575] (1.00) | PE | IL1R1 | Surface | 1: | 20 |
| 28 | YG04 [625] (1.00) | PE-CF594 | TCRgd | Surface | 1: | 20 |
| 29 | YG06 [675] (1.00) | PE-Cy5 | CD137 | Surface | 1: | 20 |
| 30 | YG07 [700] (1.00) | PE-Cy5.5 | FOXP3 | *Intracellular* | 1: | 20 |
| 31 | YG10 [780] (1.00) | PE-Cy7 | AREG | *Intracellular* | 1: | 160 |
| 32 | YG11 [810] (1.00) | PE/Fire 810 | CD40L (CD154) | Surface | 1: | 20 |
| 33 | R02 [675] (1.00) | AF 647 | Va7.2 | Surface | 1: | 80 |
| 34 | R04 [725] (1.00) | R718 | IL-17A | *Intracellular* | 1: | 20 |
| 35 | R05 [750] (1.00) | Zombie-NIR | L/D | **Live Dead** | 1: | 500 |
| 36 | R06 [780] (1.00) | APC-H7 | CD45RO | Surface | 1: | 1280 |
| 37 | R07 [810] (1.00) | APC/Fire 810 | IL-2 | *Intracellular* | 1: | 80 |

#### Symphony Panel:

| Fluorophore | Antigen | Cocktail | Dilutions | |
|---|---|---|---|---|
| BUV395 | CD8 | *Intracellular* | 1: | 80 |
| Blue, eBiosciences (UV450) | DEAD | **Live Dead** | 1: | 500 |
| BUV496 | CD3 | Surface | 1: | 40 |
| BUV563 | CD56 | Surface/*Intracellular* | 1: | 160 |
| BUV661 | CCR7 | Surface | 1: | 40 |
| BUV805 | CD45 | Surface | 1: | 80 |
| V500 | CD16 | *Intracellular* | 1: | 160 |
| BV570 | CD45RA | Surface | 1: | 80 |
| BV605 | CCR6 | Surface | 1: | 20 |
| BV650 | CD25 | Surface | 1: | 40 |
| BV711 | CD14 | Surface | 1: | 80 |
| BV750 | CD103 | Surface | 1: | 160 |
| BV785 | CD127 | Surface | 1: | 20 |
| FITC | IFNg | *Intracellular* | 1: | 20 |
| BB630 | NKp46 Streptavidin | Surface/*Intracellular* | 1: | 640 |
| BB700 | CD161 | Surface | 1: | 20 |
| PE | RORyT | *Intracellular* | 1: | 10 |
| PE-CF594 | TCRgd | Surface | 1: | 20 |
| PE-Cy5.5 | Foxp3 | *Intracellular* | 1: | 20 |
| PE-Cy7 | TNFa | *Intracellular* | 1: | 20 |
| A647 | Va7.2 | Surface | 1: | 40 |
| R718 | IL-17a | *Intracellular* | 1: | 80 |
| APC-H7 | CD4 | *Intracellular* | 1: | 40 |

#### Antibody details

| Panel | Antigen | Clone | Vendor | Catalogue # |
|---|---|---|---|---|
| 37c | CD4 | SK3 | Biolegend | 344686 |
| Symphony | CD4 | RPA-T4 | BD | 560158 |
| BOTH | CD8a | RPA-T8 | BD | 563795 |
| BOTH | CD3 | UCHT1 | BD | 612940 |
| 37c | CD161 | DX12 | BD | 748949 |
| Symphony | CD56 | NCAM16.2 | BD | 565704 |
| 37c | CD39 | TU66 | BD | 751269 |
| BOTH | CCR7 (CD197) | 2-L1-A | BD | 749824 |
| 37c | ICOS (CD278) | DX29 | BD | 749665 |
| BOTH | CD45 | HI30 | BD | 612891 |
| 37c | CD25 | 2A3 | BD | 564033 |
| Symphony | CD25 | M-A251 | BD | 563719 |
| 37c | IL-17F | O33-782 | BD | 561338 |
| 37c | CD28 | CD28.2 | BD | 566110 |

| | | | | |
|---|---|---|---|---|
| Symphony | CD16 | 3G8 | BD | 561393 |
| 37c | CD27 | M-T271 | BD | 740167 |
| BOTH | CD45RA | HI100 | Biolegend | 304132 |
| 37c | (CD279) PD1 | EH12.1 | BD | 563245 |
| Symphony | CCR6 | 11A9 | BD | 562724 |
| 37c | KI67 | B56 | BD | 563757 |
| 37c | CD69 | FN50 | BD | 563836 |
| Symphony | CD14 | MΦ P9 | BD | 563372 |
| BOTH | CD103 | Ber-ACT8 | BD | 747099 |
| BOTH | CD127 | HIL-7R-M21 | BD | 563324 |
| 37c | TIM3 (CD366) | 7D3 | BD | 565568 |
| 37c | IFNg | B27 | BD | 569265 |
| Symphony | IFNg | B27 | BD | 554700 |
| Symphony | NKp46 | 9E2 | BioLegend | 331906 |
| Symphony | Streptavidin BB630 | | BDBiosciences | 624294 |
| 37c | CTLA4 (CD152) | BNI3 | BD | 624294 |
| 37c | IL-13 | JES10-5A2 | BD | 624295 |
| 37c | IL-22 | MH22B2 | BD | 567688 |
| Symphony | CD161 | DX12 | BD | 745791 |
| 37c | Granzyme B | GB11 | BD | 570496 |
| 37c | TNFa | MAb11 | BD | 569091 |
| Symphony | TNF | Mab11 | BD | 557647 |
| 37c | IL1R1 (CD121a) | polyclonal | R&D systems | FAB269P |
| Symphony | RORyT | Q21-559 | BD | 563081 |
| BOTH | TCRgd | B1 | BD | 562511 |
| 37c | CD137 | 4B4-1 | BD | 551137 |
| BOTH | FOXP3 | PCH101 | Invitrogen | 35-4776-42 |
| 37c | AREG | AREG559 | Invitrogen | 25-5371-42 |
| 37c | CD40L (CD154) | 24-31 | Biolegend | 310857 |
| BOTH | Va7.2 | 3C10 | Biolegend | 351726 |
| BOTH | IL-17A | N49-653 | BD | 566938 |
| 37c | CD45RO | UCHL1 | BD | 561137 |
| 37c | IL-2 | MQ1-17H12 | Biolegend | 500355 |

**Supplementary Table S3: Lung donor characteristics**

**Deceased donor cohort**

| Characteristics | Donor cohort (N=11) |
|---|---|
| Age (median, interquartile range [IQR]) | 56 (6-59) |
| Female sex | 4 (36%) |
| Cause of death | |
| Head trauma | 5 (46%) |
| Stroke | 4 (36%) |
| Anoxic brain injury | 2 (18%) |
| Baseline immunosuppression | 0 |
| Days of mechanical ventilation (median, IQR) | 5 (4-6) |

**ILD donor cohort**

| Characteristics | ILD cohort (N=10) |
|---|---|
| Age (median, interquartile range [IQR]) | 58 (52-65) |
| Female sex | 4 (40%) |
| ILD-related diagnosis | |
| Idiopathic interstitial pneumonia | 4 (40%) |
| Rheumatologic-associated disease | 2 (20%) |
| Idiopathic pulmonary fibrosis | 2 (20%) |
| Other | 2 (20%) |
| Baseline prednisone (%) | 4 (40%) |
